# Adeno-Associated Virus 2 (AAV2) - induced RPA exhaustion generates cellular DNA damage and restricts viral gene expression

**DOI:** 10.1101/2025.04.22.649920

**Authors:** Monnette F. Summers, MegAnn K. Haubold, Marcel Morgenstern, Phoenix Shepherd, Clairine I.S. Larsen, Ava E. Bartz, Gopishankar Thirumoorthy, Robert N. Kirchdoerfer, Joshua J. Coon, Kavi P. Mehta, Kinjal Majumder

**Affiliations:** Institute for Molecular Virology, University of Wisconsin-Madison, Madison, WI, 53706; McArdle Laboratory for Cancer Research, School of Medicine and Public health, Madison, WI, 53706; Department of Biomolecular Chemistry, School of Medicine and Public health, Madison, WI, 53706; National Center for Quantitative Biology of Complex Systems, University of Wisconsin-Madison, Madison, WI, 53706; Department of Biochemistry, University of Wisconsin-Madison, Madison, WI, 53706; Center for Quantitative Cell Imaging, University of Wisconsin-Madison, Madison, WI, 53706; Department of Comparative Biosciences, School of Veterinary Medicine, Madison, WI, 53706; Department of Chemistry, University of Wisconsin-Madison, Madison, WI, 53706; Morgridge Institute for Research, Madison, WI, 53715; University of Wisconsin Carbone Cancer Center, Madison, WI, 53712

**Keywords:** parvovirus, DNA damage response, Adeno-Associated Virus, replication stress, gene therapy, recombinant AAV2 gene therapy

## Abstract

Parvoviruses are single-stranded DNA viruses that have been modified to serve as vehicles for therapeutic transgene delivery in the form of recombinant Adeno-Associated Virus (rAAV2) vectors or rodent parvovirus-derived oncolytic agents. Infection with viruses of the *Parvoviridae* family induces a cellular DNA Damage Response (DDR) signal that supports virus replication. However, it remains unknown whether rAAV2 vectors or non-replicative AAV2 genomes induce cellular DDR signals, which might be deleterious to the cell. To determine the impact of AAV2/rAAV2 genomes on the integrity of the host chromosome, we have pulsed AAV2/rAAV2 infected cells with BrdU analogs followed by single-molecule imaging of the cellular replisomes and proteomic analysis of the host replication forks. We discovered that non-replicative AAV2/rAAV2 genomes are sufficient to induce replication stress on the host genome, leading to DDR signals in a dose-dependent manner. Moreover, infection with replication-competent AAV2 leads to enrichment of replication stress proteins, DNA repair factors and RNA processing machinery on cellular replication forks. However, neither the AAV2 Inverted Terminal Repeats (ITRs) that are retained in rAAV2s nor empty capsids are sufficient to induce host-cell replication stress. Strikingly, incoming AAV2 genomes associate with the single-stranded DNA binding protein RPA in host cells in a dose-dependent manner, progressively shortening cellular replication forks. These elevated levels of AAV2-induced cellular replication stress eventually leads to accumulation of DDR signals in the nucleus. Chemical inhibition of RPA activity and RNAi-mediated knockdown leads to de-repression of the AAV2 genome, increasing Rep 68/78 gene expression. Ectopic expression of RPA rescues AAV2-induced replication stress. Taken together, our findings suggest that depletion of cellular stores of RPA molecules by competing AAV2 genomes restrict viral gene expression and cause cellular DNA damage.

**AUTHOR SUMMARY:** Adeno-Associated Viruses 2 (AAV2) are modified to design therapeutic gene therapy vectors, but how they interact with the guardians of host DNA remains unknown. In this work, we show that AAV2 genomes compete with the host cell for the single-stranded DNA binding protein RPA, rendering the host vulnerable to replication stress leading to both suppression of the viral gene expression and induction of cellular DNA breaks. These findings provide insights into how gene therapies delivered at high doses could have genotoxic effects, underscoring the importance of engineering AAV2-based gene therapy platforms that express efficiently at lower doses.

## INTRODUCTION

DNA breaks generated by viral and non-viral agents activate DNA Damage Response (DDR) signals that have evolved to ensure the fidelity of the genetic code. These signals facilitate DNA repair by recruiting cellular DNA processing factors that shut down transcription in the vicinity of the DNA break, process the DNA strand and restore the integrity of the genome. Primarily regulated by the evolutionarily conserved Phosphatidylinositol-3-Kinase-like kinases ATM, ATR and DNAPK, these DDR signals are usurped by viruses to regulate the outcome of viral infection. To carry out their life cycles in the nuclear compartment, DNA viruses either trigger (parvoviruses [1, 2], polyomaviruses [3, 4] and papillomaviruses [5, 6]) or inactivate (adenovirus [7–11] and herpesviruses [12]) cellular DDR signaling pathways for distinct infectious outcomes (such as the establishment of latency [12] or production of progeny virus [13–15]). DNA viruses interfere with DDR signaling pathways activated by ATM, ATR or DNA-PK signals using their viral proteins (adenovirus) or nucleic acids (herpesviruses and adenoviruses). Parvoviruses are single-stranded DNA viruses that can replicate in cells in S-phase (referred to as autonomous parvovirus) or with the help of co-infecting viruses (known as dependoparvovirus). Replication of the dependoparvovirus Adeno-Associated Virus Type 2 (AAV2) induces a cellular DDR signal using the DNA-PK pathway [16], but the signals associated with non-replicative (lacking a co-infecting virus; hereafter referred to as mono-infection) AAV2 infection remain largely unknown.

The cellular ATR signaling pathway has evolved to respond to single-stranded DNA breaks in the nucleus that are generated by replication stress and transcription-replication conflicts [17, 18]. Viruses deploy a diverse set of programs to dysregulate ATR-mediated signals for their benefit (reviewed in [17]). Rift Valley Fever Virus and the autonomous parvovirus Minute Virus of Mice (MVM), for example, express proteins that inactivate components of the ATR pathway [19, 20]. Papillomavirus E2 proteins and Polyomavirus Large-T antigen interact with ATR-pathway proteins (TOPBP1 [21] and Claspin [22] respectively) and components of eukaryotic replisomes to control ATR-dependent signals. The genomes of the autonomous parvovirus Minute Virus of Mice (MVM) sequester the cellular stores of Replication Protein A (RPA), the principal sensor of single stranded DNA in the nucleus, which in turn initiates ATR signaling, thereby rendering the host genome vulnerable to replication stress. This culminates in the generation of extensive cellular DNA damage [23], contributing to the induction of a potent pre-mitotic cell cycle arrest at the G2/M border [24], an outcome that has been leveraged to engineer oncolytic virotherapies using protoparvoviruses [25]. Since MVM replication rapidly generates single-stranded DNA, double stranded DNA and RNA-DNA hybrid intermediates [26], it remains unknown what aspect of viral replication activates cellular DDR signals. In the absence of co-infecting “helper” viruses, non-replicative Adeno-Associated Viruses type 2 (AAV2) and the recombinant AAV2 (rAAV2) gene therapy vectors that are derived from them represent a tractable system to dissect the DDR signals activated by viral infection [10].

We have previously discovered that genomes of autonomous parvoviruses like MVM and dependoparvoviruses like AAV2 localize to cellular sites of DNA damage [2, 27]. At these sites, MVM forms replication centers called Autonomous Parvovirus Associated Replication (APAR) bodies [28, 29], whereas AAV2/rAAV2 genomes form subnuclear structures where their genomes are converted from single-stranded to double-stranded forms [30]. MVM-generated subnuclear structures are formed in proximity to large host chromatin domains spanning multiple megabases [2, 31] but AAV2-associated nuclear structures associate with distinct regions that colocalize with sharp chromatin sites that are in the vicinity of DNA breaks [27]. Many of these cellular DDR sites are fragile genomic regions [32–34] where the cellular replication and transcription machineries collide to form transient secondary structures that recruit DNA processing factors (such as ATR), serving as fertile milieu for establishment and replication of viruses in their vicinity [35–38]. ATR-mediated signals regulate the localization to cellular DDR sites of the parvovirus non-structural protein NS1 (for MVM) and Rep 67/78 (for AAV2) and are essential for efficient virus replication [27, 39]. At late stages of viral life cycle, MVM inactivates the ATR signaling pathway by sequestering Casein Kinase 2 (CK2), blocking its ability to phosphorylate downstream substrates [40]. However, it remains unknown whether ATR signaling is altered during AAV2 mono-infection.

The outcome of AAV2 mono-infection on host DDR pathways remains controversial. Some of these prior studies posit that AAV2 genomes persist in the nuclear compartment as a molecular mimic of stalled host replication forks [41], leading to local activation of CHK1-related signals (downstream of ATR activation). These CHK1-dependent signals cause cell death in cells that lack p53 signaling [42, 43]. On the other hand, cellular DDR signals associated with double stranded breaks monitored by phosphorylation of H2AX (referred to as γH2AX) are not always detected in AAV2/rAAV2-infected host cells [44, 45]. In contrast, transduction of human Embryonic Stem (ES) cells with rAAV2 leads to host genome instability and p53-dependent cell death [46]. Nevertheless, it is incontrovertible that parvoviruses, lacking polymerases of their own, rely exclusively on the host for polymerases to amplify their genomes. Cellular DDR signaling pathways regulate the licensing of replicative DNA polymerases alpha, delta and epsilon [47–49]; as well as translesion polymerases eta, zeta and kappa [50]. While inhibition of DNA polymerases alpha and delta inhibits the replication of Parvovirus B19V [51], eta and kappa knockdowns decrease Human Bocaparvovirus (HBoV1) replication [52]. Though the absence of polymerase eta and kappa reduces AAV2 replication [44], this occurs only in the presence of adenovirus “helper” proteins (E1, E2, E4orf6 and VA-RNA [53, 54]). Importantly, inhibition of ATR signaling arrests AAV2 replication [44], possibly because ATR regulates polymerase eta and kappa function [55, 56]. Since repair polymerases must be licensed by host activation signals [57], these observations additionally suggest that AAV2 must activate local signals that are yet to be elucidated. However, AAV2 monoinfection (i.e. infection in the absence of a co-infecting helpervirus) does not support viral genome amplification, eliminating the need of cellular DDR signals to license virus replication, suggesting that cellular DDR signals are generated as a host response to viral infection and not as a means of viral pathogenesis.

In this study, we have investigated whether and how AAV2 mono-infection impacts the stability of the eukaryotic replisome using assays that label the nascent host DNA strand. Imaging of the eukaryotic replisomes and mass-spectrometry-based identification of the proteins associated with replicating DNA revealed an enrichment of replication-stress markers upon AAV2 infection. This is caused by the AAV2 genomes serving as molecular decoys for the single-stranded DNA binding protein RPA, rendering the host genome vulnerable to replication stress. Surprisingly, while AAV2-induced RPA exhaustion leads to cellular DNA damage at high doses, the binding of RPA molecules to AAV2 restricts viral gene expression. Taken together, our findings define how AAV2 genomes modulate the replication fork stability in host cells that can cause genomic DNA damage, explaining how viral genes are repressed by replication stress proteins in the absence of coinfecting helperviruses.

## RESULTS

### Replicative and non-replicative AAV2 genomes induce replication stress in cycling cells

Parvovirus replication induces cellular DDR signals monitored by increase in γH2AX levels and comet tails associated with DNA fragmentation [1, 2, 44], but it remains unclear whether mono-infection induces DNA damage. Consistent with this model, AAV2 replication in the presence of helper viruses (Adenovirus or Herpesvirus) or ectopically expressed Adenovirus helper proteins induces a robust cellular DDR but the impact of AAV2 mono-infection on the cellular DDR pathways is much less pronounced. Since the AAV2 genome mimics a stalled-fork [41], we hypothesized that these entities competes with the host genome for cellular replication-stress factors. To determine whether AAV2 mono-infection impacts the stability of host cell replication forks, we utilized single-molecule DNA Fiber Assays (DFAs; [58]). DFAs measure the length of host cell replication tracts using sequential pulses of the halogenated nucleoside analogs Iododeoxyuridine (IdU) and Chlorodeoxyuridine (CldU) followed by spreading of DNA fibers on positively charged slides analyzed by confocal imaging. As schematized in Fig. 1A, DFAs of AAV2-infected HEK293T cells at 24 hours post-infection (hpi) revealed a significant shortening of cellular replication forks (representative examples in Fig. 1B) during AAV2 mono-infection from CldU lengths of 4.1 μm in Mock-infected cells to 3.1 μm in AAV2-infected cells. Interestingly, cells transfected with only the pHelper plasmid (expressing the adenovirus helper proteins E2A, E4orf6 and VA-RNA) are sufficient to shorten host-cell replication forks (measured by both IdU and CldU labels, Fig. 1C, 1D). These observations suggested that Adenovirus proteins that degrade cellular DDR factors like MRE11 exacerbate cellular replication stress (as previously observed and published, [11]). To validate these observations in an independent cell line model using orthogonal techniques, we performed AAV2 infection of U2OS osteosarcoma cells and monitored the ability of host replisomes to incorporate 5-Ethynyl-2’-deoxyuridine (EdU) analogs by conjugating fluorophores using Click-iT chemistry. Infection of U2OS cells with AAV2 for 24 hours at a multiplicity of 5,000 viral genomes per cell led to fewer and smaller EdU-positive foci (representative image in Fig. 1E and quantified in Fig. 1F). The median number of EdU-positive foci in AAV2-infected cells decreased to 3 from 6 in mock infected cells (Fig. 1F), phenocopying our DFA analysis in 293T cells. To determine when AAV2 mono-infection induces cellular replication stress, we performed DFAs at 12 hpi, 24 hpi and 36 hpi. As shown in Fig. 1G (IdU) and 1H (CldU), there was no robust change in median fork lengths from the onset of infection to 12 hpi, indicating that AAV2-induced replication stress is not an acute response. However, there was a substantial decrease in median fiber lengths between 12 hpi and 24 hpi, which plateaued by 36 hpi (Fig. 1G, 1H). This plateau of AAV2-induced replication fork shortening at 36 hpi suggested that the host cell might deploy mechanisms to overcome the virus-induced replication stress. Taken together, our observations suggested that AAV2 mono-infection induces replication stress on the host genome.

**Fig. 1:**
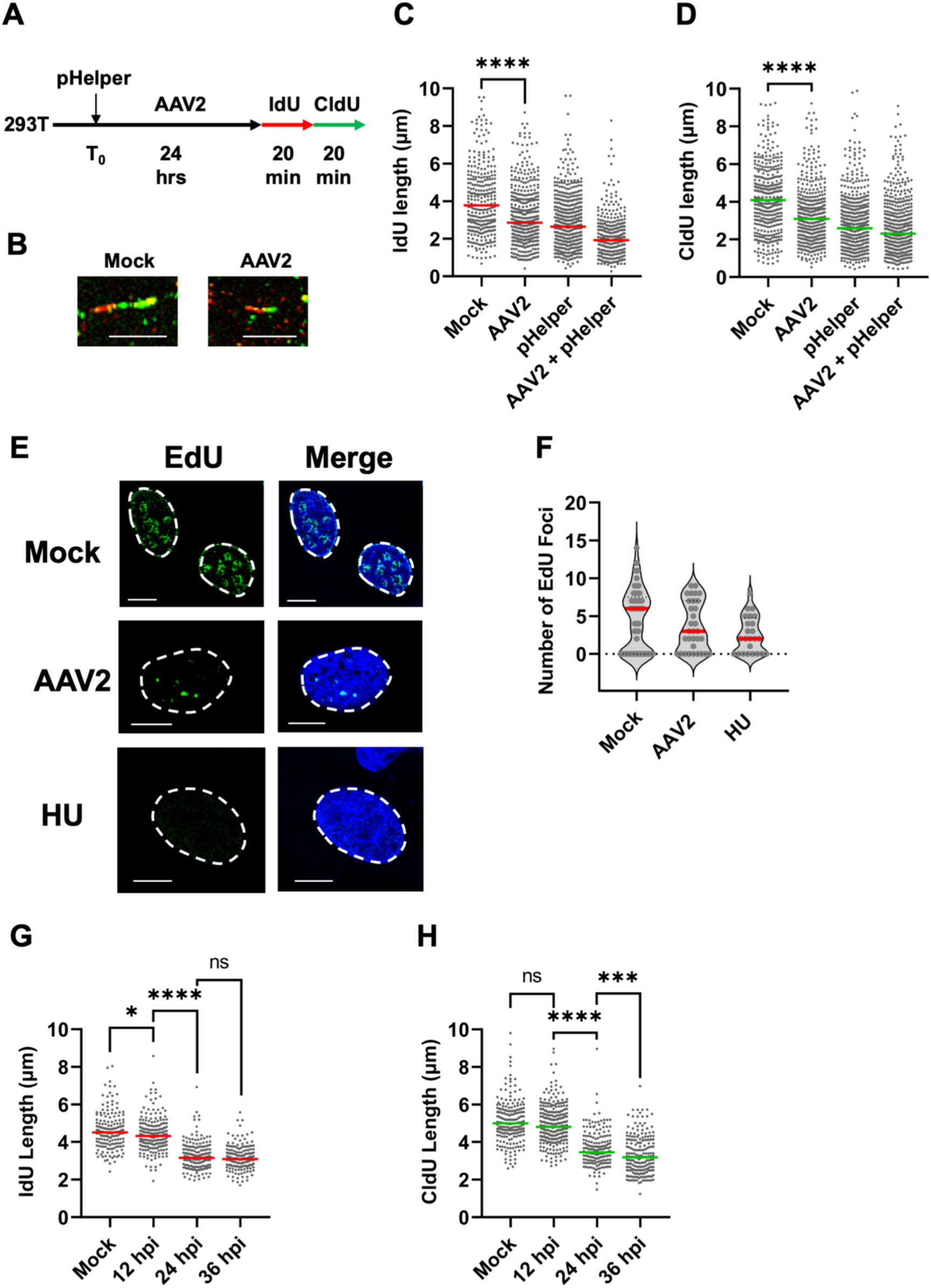
Replicative and non-replicative AAV2 genomes induce replication stress in cycling cells. (A) Schematic of the timeline of AAV2 infection of 293T cells followed by sequential pulsing with IdU and CldU for DNA fiber analysis. (B) Representative images of single DNA fibers in mock (left) compared with AAV2-infected 293T cells (right) at 24 hpi. The red portions of the fibers represent IdU label and green portions represent CldU labels. The horizontal scale bar represents 5 micrometers. (C, D) DNA fiber length measurements were indicated by each datapoint for (C) IdU-labelled fibers and (D) CldU-labelled fibers. The median lengths of many measurements of IdU and CldU labelled fibers are represented by red and green bars respectively. At least 150 individual fibers were measured for each condition for three independent replicates of AAV2 infections of 293T cells. Statistical significance was measured by Mann Whitney Wilcoxon test, **** represents P < 0.0001. (E) Representative images of cellular replication forks (green) in uninfected U2OS cells compared with U2OS cells infected with AAV2 and treated with 2μM hydroxyurea (HU) for 12 hours. Cells were pulsed with EdU followed by conjugation of Alexa Fluor 488 using click chemistry. The cell nucleus is stained with DAPI (blue) and the nuclear borders are demarcated with a white dashed line. The scale bar represents 5 micrometers. (F) Quantification of EdU labelling in 1E with each measurement represented by a datapoint on the graph. The median values are represented by red horizontal bars from EdU foci counted in over 30 U2OS cell nuclei. (G, H) Measurement of single molecule DNA fiber lengths in AAV2 infected 293T cells at the indicated timepoints of infection which were followed by sequential pulses of IdU and CldU for 20 minutes each. The data is presented as IdU lengths (G) and CldU lengths (H) with the red and green horizontal bars showing the respective median values. Statistical significance was measured by Mann Whitney Wilcoxon test, **** represents P < 0.0001, *** P < 0.001, * P < 0.05, ns represents differences that are not statistically significant.

### Recombinant AAV2 vector genomes induce replication stress in host cells

Recombinant AAV2 gene therapy vectors have been engineered from AAV2 by replacing the Rep-Cap containing open reading frames with therapeutic transgenes that are the under control of transcriptional regulatory elements [59]. Transduction of cells with rAAV2 vectors have previously yielded conflicting results regarding DNA damage induction, with early reports indicating that rAAV2 vectors activate CHK1 kinase globally [42], induce cell death in transformed cycling cells [43] and toxicity in human Embryonic Stem (ES) cells [46]. Surprisingly, follow-up studies found that non-replicative AAV2 and rAAV2 are insufficient to provoke global DDR signals [16, 44, 60]. To determine whether rAAV2 genomes disturb the cellular replisome, we performed DFAs in 293T cells transduced with rAAV2 at an MOI of 5,000 viral genomes per cell for 24 hours (Fig. 2A). We compared the cellular replisome in rAAV2-transduced cells with that of self-complementary AAV2 (scAAV2) that contain only one AAV2-ITR and form a double-stranded DNA molecule soon after nuclear entry (removing the need for scAAV2s to undergo DNA processing). As shown in Fig. 2B and Fig. 2C, both rAAV2 and scAAV2 transduction led to a shortening of cellular replication forks measured by shortening in the median lengths of both nucleoside analog (IdU and CldU). The median lengths of cellular replication forks in AAV2-infected cells were comparable to that of rAAV2/scAAV2-transduced (lacking the Rep and Cap ORFs) 293T cells at 24 hpi. This observation indicated that the non-coding components of AAV2 genomes might be involved in replication stress and the low levels of Rep 68/78 protein generated by AAV2 mono-infection is not sufficient to account for the observed cellular replication stress. The similar levels of replication stress induced by AAV2, rAAV2 and scAAV2 suggested that either be the ITR (which is shared between at three genomic forms) or the viral particle (VP; also shared between all three infection/transductions) might be sufficient to contribute to AAV2-induced replication fork shortening.

**Fig. 2:**
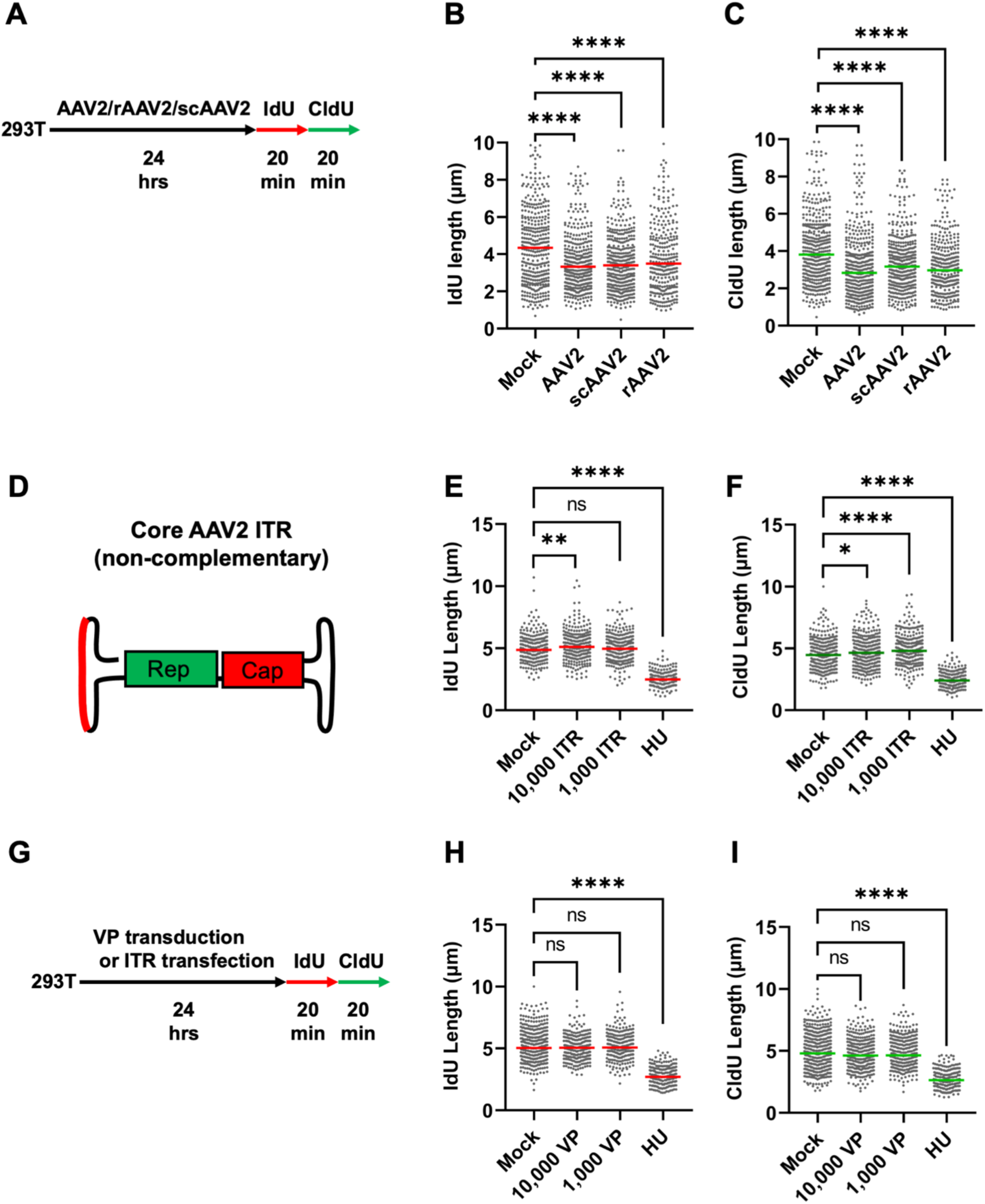
Recombinant AAV2 vector genomes induce replication stress in host cells. Schematic of the timeline of AAV2 infection and rAAV2/scAAV2 transduction of 293T cells followed by sequential pulsing with IdU and CldU for 20 minutes each prior to DNA fiber analysis. (B, C) DNA fiber length measurements were indicated by each datapoint for (B) IdU-labelled fibers and (C) CldU-labelled fibers. The median lengths of many measurements of IdU and CldU labelled fibers in AAV2 infected cells and rAAV2/scAAV2 transduced cells are represented by red and green bars respectively. At least 150 individual fibers were measured for each condition for three independent replicates of AAV2 infections of 293T cells. Statistical significance was measured by Mann Whitney Wilcoxon test, **** P < 0.0001, *** P < 0.001, ** < 0.01, ns represents differences that are not statistically significant. (D) Schematic of the AAV2 genome with red line indicating the non self-complementary region of the ITR transfected into 293T cells for 24 hours prior to sequential IdU/CldU labelling for 20 mins each followed by DNA fiber analysis. (E, F) DNA fiber length measurements were indicated by each datapoint for (E) IdU-labelled fibers and (F) CldU-labelled fibers. The median lengths of many measurements of IdU and CldU labelled fibers in empty VLP transduced cells are represented by red and green bars respectively. (G) Schematic of the timeline of empty assembled VLP transduction or ITR element transfection of 293T cells followed by sequential pulsing with IdU and CldU for 20 minutes each prior to DNA fiber analysis. (H,I) Representation of the lengths of the (H) IdU and (I) CldU forks with median lengths represented by red and green bars respectively. At least 150 individual fibers were measured for each condition for three independent replicates of AAV2 infections of 293T cells. Statistical significance was measured by Mann Whitney Wilcoxon test, **** represents P < 0.0001, ns represents differences that are not statistically significant.

### Empty AAV2 capsids and ITR elements are not sufficient to induce host-cell replication stress

To determine whether the AAV2 ITR element is sufficient to induce host-cell replication stress, we transfected 293T cells with 10,000 and 1,000 copies of a non-self-complementary ITR sequence containing oligonucleotides for 24 hours before pulsing cells with IdU/CldU for DFAs (Fig. 2D). We compared the lengths of the replication forks in these cells with that of cells transfected with equivalent number of copies of pUC18 plasmid DNA and cells treated with HU. Measurement of the replication forks in these cells revealed that the ITR elements were not sufficient to cause cellular replication stress measured by IdU (Fig. 2E) or CldU lengths (Fig. 2F). To determine whether the AAV2 Capsid is sufficient to induce cellular replication stress, we transduced 293T cells with 10,000 or 1,000 copies of the empty AAV2 capsids for 24 hours before performing DFAs (according to the schematic in Fig. 2G). As shown in Fig. 2H (IdU) and 2I (CldU), transduction of these empty capsids was not sufficient to shorten the host-cell replication fibers. These findings suggested that the non-self-complementary ITR region and empty viral particles are not sufficient to induce cellular replication stress.

### Characterization of the AAV2-induced alterations to the human replisome using iPOND-MS

To determine which host proteins are altered at eukaryotic replisomes by AAV2 infection and corroborate our observations of AAV2-induced replication stress (using DFAs and EdU labelling), we utilized the proteomics approach iPOND [Isolation of Proteins On Nascent DNA; [61]]. iPOND uses EdU-based labelling followed by Click-chemistry-mediated biotinylation coupled with streptavidin pulldowns to identify the proteins associated with nascent DNA[61]. We performed iPOND analysis in 293T and U2OS cells infected with AAV2 at an MOI of 10,000 vg/cell at 24 hpi. As shown in Fig. 3A, iPOND-MS analysis of AAV2-infected cells yielded 1084 proteins that were dysregulated in U2OS cells and 1606 proteins in 293T cells. Out of these, 822 proteins were shared between U2OS and 293T cells (Fig. 3A). Amongst the shared proteins that are dysregulated by AAV2, 414 were enriched and 408 were depleted at host-cell replication forks (Fig. 3B). Unbiased pathway analysis of these 822 proteins revealed that these factors are involved in DNA replication, double-stranded DNA break repair and chromosome organization (Fig. 3C). Strikingly, several proteins are also involved in RNA processing, RNA localization and splicing (Fig. 3C). Host proteins enriched at cellular replication forks during infection included DNA processing factors such as GINS, TOP2A, PRIM1/2, RPA2 and XRCC1; DDR signaling proteins such as CSNK2A1/3; cell cycle regulator proteins such as TFDP2 and SFN1; as expected for cells undergoing replication stress. Host DDR-associated proteins depleted at cellular replication forks in AAV2-infected cells included SIK3, DAXX and NBN (Fig. 3D). To determine whether these host replication fork-associated proteins regulate AAV2 gene expression, we performed RNAi-knockdowns in 293T cells, infected with AAV2 for 24 hpi and assessed their impact on Rep 68/78 transcript levels. As shown in Fig. 3E, knockdowns of XRCC1, SIK, PRIM1 and TFDP2 did not impact AAV2 gene expression substantially. However, knock-down of RPA2 (also known as RPA32) led to a substantial de-repression of AAV2 gene expression, causing a 10-fold increase of Rep 68/78 transcripts (Fig. 3E, left y-axis). Strikingly, absence of CSNK2A1/3 led to a 300-fold increase in AAV2 gene expression relative to Mock knockdown cells (Fig. 3E, right y-axis). These observations corroborate and build on the recent findings of CSNK1 serving as a host restriction factor for scAAV2 transduction [62]. These observations orthogonally validated the cellular replication stress monitored by DFAs and EdU-labelling, additionally identifying host factors that might regulated changes at the cellular replisomes.

**Fig. 3:**
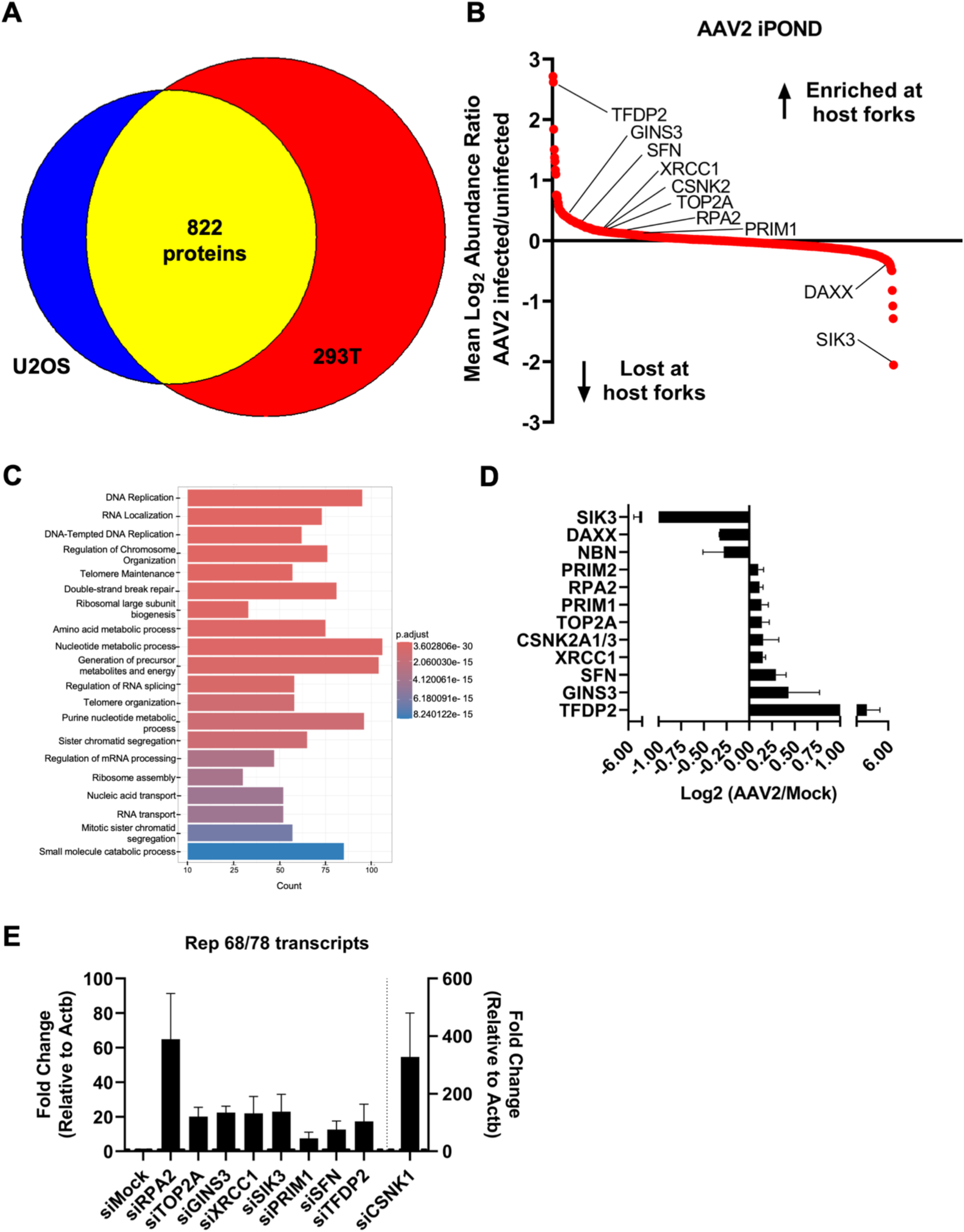
Characterization of the AAV2-induced alterations to the human replisome using iPOND-MS. (A) Venn diagram depicting the number of detected proteins enriched in 293T cells (red), U2OS cells (blue) and shared between the two cell types (yellow). (B) Scatter plot comparing the mean log2 abundance ratios between AAV2 infected and uninfected cells among the 822 proteins identified in the datasets in (A) to its corresponding rank in decreasing order. (C) Gene ontology analysis of proteins enriched on nascent DNA in AAV2-infected 293T cells indicating the pathways that are associated with these proteins. (D) Comparison of protein enrichment on nascent DNA between Mock infected and AAV2-infected cells represented as mean of log2 values between U2OS and 293T cells focused on proteins indicated in Fig. 3B. (E) RT-qPCR analysis of the Rep 67/78 transcript levels in AAV2-infected 293T cells at 24 hours post-infection that were transfected with the indicated siRNAs. The AAV2 transcripts are compared with Beta-actin transcripts as loading control and siMock-transfected cells set to one to calculate fold-change. Data is represented as mean ± SEM of from two independent replicates of RNAi knockdowns followed by infections. Right y axis denotes the fold change relative to beta-actin in siCSKN1 transfected cells.

### AAV2 genomes associate with cellular RPA and RPA overexpression rescues AAV2-induced replication stress

Since cellular RPA2 is enriched on the host replisome and its knockdown derepressed Rep 68/78 transcripts, we hypothesized that RPA2 plays a dual role at the AAV2-host replisome interface. Consistent with this, we have previously discovered that genomes of the related autonomous parvovirus MVM induces cellular DDRs by depleting the host cell of RPA2 molecules to exacerbate replication stress [23]. However, since AAV2 genomes do not replicate in the absence of helpervirus co-infection, they are unlikely to cycle between single- and double-stranded intermediates that characterize parvovirus genome amplification [26, 63, 64]. To determine whether non-replicative AAV2 genomes associate with components of the heterotrimeric RPA complex (made up of RPA14, RPA32 and RPA70 [65]), we performed ChIP-qPCR assays for these individual RPA subunits on the 5’ end of the AAV2 genome. Interestingly, all three of the RPA subunits were bound to AAV2 genomes in the nucleus at 24 hpi (Fig. 4A). To examine whether assembly of the entire RPA heterotrimer on the AAV2 genome leads to RPA activation, and if this depends on the number of AAV2 genome copies, we performed ChIP-qPCR for RPA32 phosphorylated at Serine 8 (which regulates the activation of replication checkpoint signals [66]) at increasing MOI’s of AAV2 genomes in 293T cells at 24 hpi (Fig. 4B). We discovered that high MOIs of AAV2 in 293T cells associated with phospho-RPA32, with a stepwise increase in interactions from 10,000 vg/cell to 20,000 vg/cell (Fig. 4B). We confirmed that RPA32 phosphorylation is inhibited on the AAV2 genome using TDRL-505 [67], an allosteric inhibitor of RPA32’s phosphorylation (Fig. 4C). While inhibition of RPA32 phosphorylation by TDRL-505 (using the schematic illustrated in Fig. 4D) was sufficient to shorten cellular replication forks monitored by IdU and CldU lengths (iRPA samples in Fig. 4E, 4F), these defects on the host replication fork was not enhanced further in the presence of AAV2 infection (iRPA+AAV2 in Fig. 4E, 4F). This suggests that the ability of RPA32 to bind the AAV2 genome or host replication forks plays a key role in protecting the genome from replication stress. Strikingly though, inhibition of RPA32 activation led to derepression of the AAV2 genome, increasing the levels of Rep 68/78 transcripts generated in the 293T cells (Fig. 4G, left) and U2OS cells (Fig. 4G, right). However, RPA32 inhibition did not impact the intensity of rAAV2 transducing a GFP transgene (Fig. 4H). These findings suggested that RPA32-mediated regulation of AAV2 gene expression is distinct from that of rAAV2. To overcome the impact of RPA-exhaustion induced by AAV2 genomes, we transfected plasmids expressing the RPA subunits (pRPA) into 293T cells prior to AAV2 infection as schematized in Fig. 4D. Measurement of DNA fibers of these cells after AAV2 infection for 24 hours showed that pRPA transfection rescued AAV2-induced replication stress, elongating host replication fibers to higher than wild-type levels (IdU, Fig. 4I) or approximately equal to wild-type lengths (CldU, Fig. 4J).

**Fig. 4:**
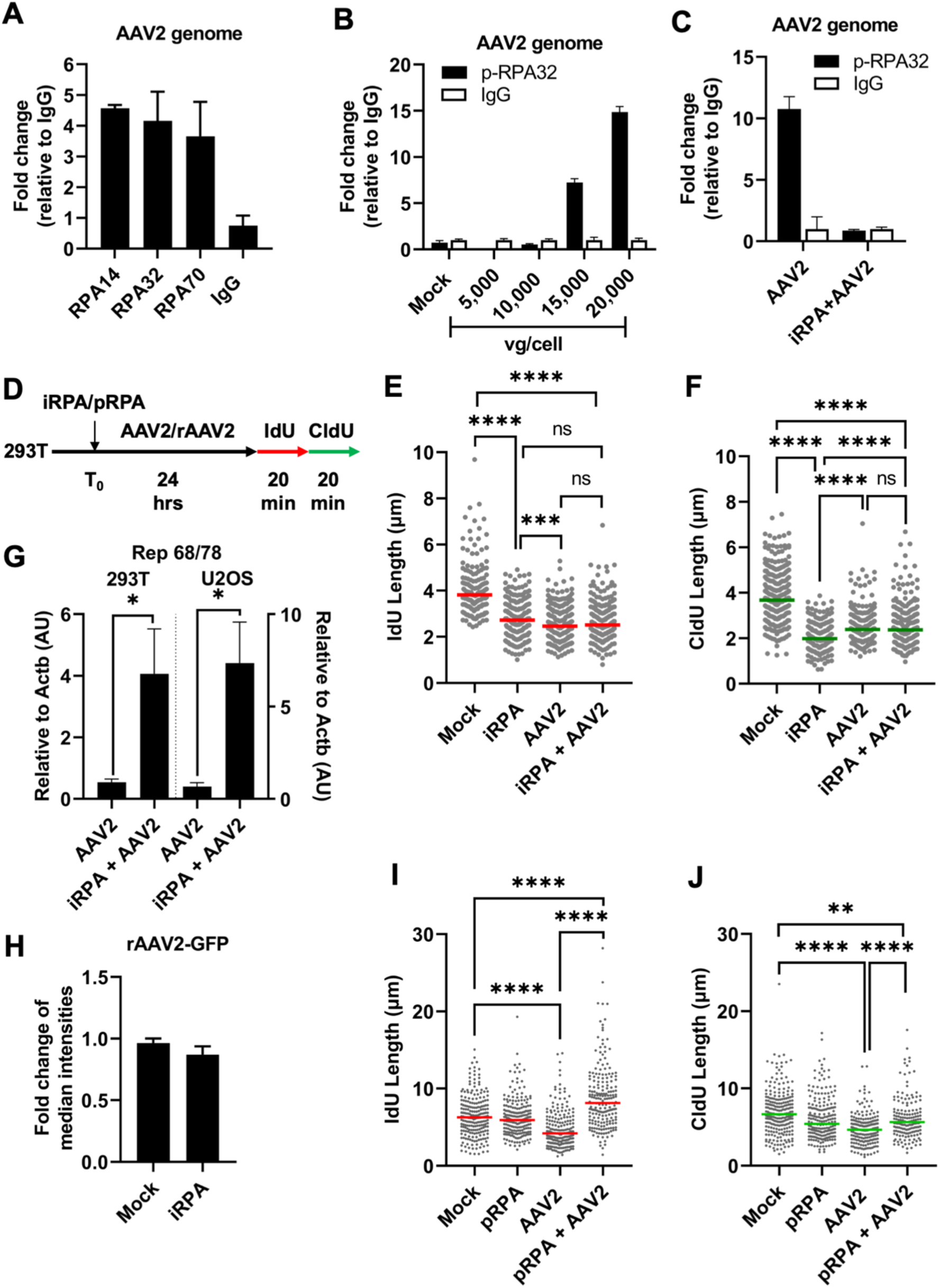
AAV2 genomes associate with cellular RPA and RPA overexpression rescues AAV2-induced replication stress. (A) Fold change of the binding of the indicated RPA subunits compared with isotype IgG to the AAV2 genome in 293T cells infected with AAV2 at an MOI of 10,000 vg/cell. Data is represented as mean ± SEM of fold change. (B) ChIP-qPCR analysis of phosphorylated RPA 32 binding to the AAV2 genome at 24 hours post-infection at the indicated MOIs compared with IgG control. (C) Impact of RPA inhibition with TDRL on phosphorylated RPA binding to the AAV2 genome at 24 hpi in 293T cells infected at an MOI of 10,000 vg/cell. Background levels of the pulldown were evaluated using IgG as control. (D) Schematic of the timeline of AAV2 infection at an MOI of 10,000 vg/cell upon being pulsed with the RPA inhibitor TDRL or pRPA transfection in 293T cells for 24 hours before sequential pulsing with IdU and CldU for 20 minutes each prior to DNA fiber analysis. (E, F) DNA fiber length measurements were indicated by each datapoint for (E) IdU-labelled DNA fibers and (F) CldU-labelled DNA fibers. The median lengths of many measurements of IdU and CldU labelled fibers in AAV2 infected cells are represented by red bar and green bar respectively. At least 150 individual fibers were measured for each condition for three independent replicates of AAV2 infections of 293T cells. Statistical significance was measured by Mann Whitney Wilcoxon test, **** represents P < 0.0001, ns represents differences that are not statistically significant. (G) AAV2 gene expression was monitored by RT-qPCR for Rep 68/78 transcripts compared with cellular *Actb* levels in AAV2 infections of 293T cells (left) and U2OS cells (right) in the presence of the RPA inhibitor TDRL-505 schematized in (D). Data presented as mean ± SEM of Rep 68/78 transcripts relative to *Actb* from at least three independent experiments. (H) Quantification of the mean fluorescence intensity of 293T cells treated with TDRL-505 prior to rAAV-GFP transduction according to the schematic in Fig. 4D. Data is represented as fold change of fluorescence intensity relative to Mock-treated cells. DNA fiber length measurements indicated by each datapoint for (I) IdU-labelled DNA fibers and (J) CldU-labelled DNA fibers in AAV2 infected cells in the presence of transfected pUC18/pRPA represented by red bar and green bar respectively as schematized in 4D. At least 150 individual fibers were measured for each condition for three independent replicates of AAV2 infections of 293T cells. Statistical significance was measured by Mann Whitney Wilcoxon test, **** represents P < 0.0001, ns represents differences that are not statistically significant.

### AAV2-mediated RPA32 exhaustion induces cellular DNA damage

To determine how competition for RPA32 molecules between host and the virus genomes impact cellular replication forks, we infected 293T cells for 24 hours with increasing the viral genomes per cell before performing DFAs by sequential IdU/CldU pulses (schematized in Fig. 5A). As shown in Fig. 5B for IdU lengths and Fig. 5C for CldU lengths, increase of MOIs of AAV2 genomes infecting 293T cells led to a dose-dependent decrease in cellular DNA synthesis. To determine whether high levels of replication stress was associated with cellular DNA damage, we labelled AAV2-infected U2OS cells (at a high MOI of 10,000 vg/cell at 24 hours) with EdU for 20 minutes, co-stained for γH2AX before performing confocal imaging. EdU labelled sites in mock-infected cells did not associate with γH2AX foci (Fig. 5D, top panel). Strikingly however, in AAV2-infected cells, the EdU labelled sites (green) colocalized with γH2AX signals (red; Fig. 5D, bottom). To determine whether induction of dose-dependent replication stress by AAV2 leads to cellular DNA damage, we measured the γH2AX foci in EdU-labelled U2OS cells infected with AAV2 at different MOIs. As shown in Fig. 5E, increasing the AAV2 genomes per cell led to an increase of γH2AX foci per cell in EdU-positive cells. To confirm the dose-dependent induction of cellular DNA damage signals, we performed western blots for γH2AX in 293T cells infected with AAV2 at progressively increasing doses, showing that γH2AX levels were enhanced in cells infected with AAV2 at MOI’s of 10,000, 15,000 and 20,000 viral genomes per cell (Fig. 5F, lanes 3-5). We validated these observations in Normal Human Dermal Fibroblasts (NHDF’s) that have low basal levels of γH2AX (Fig. 5G, lane 1), discovering that AAV2 genomes at high doses induce cellular γH2AX (Fig. 5G, lane 3). To determine where these AAV2-induced DDR signals arising on the viral genome, we performed Immuno-FISH imaging of AAV2 genomes (green) with the phosphorylated versions of RPA32, NBN (also known as NBS1) and MDC1. As shown in Fig. 5H, all the phosphorylated versions of these DDR proteins colocalized with AAV2 genomes in the nucleus of infected U2OS cells. These findings suggested that the increase of AAV2 genomes in host-cell nuclei generate a shortening of cellular replication forks by inducing RPA exhaustion that can lead to DNA damage on the cellular genome in the vicinity of AAV2.

**Fig. 5:**
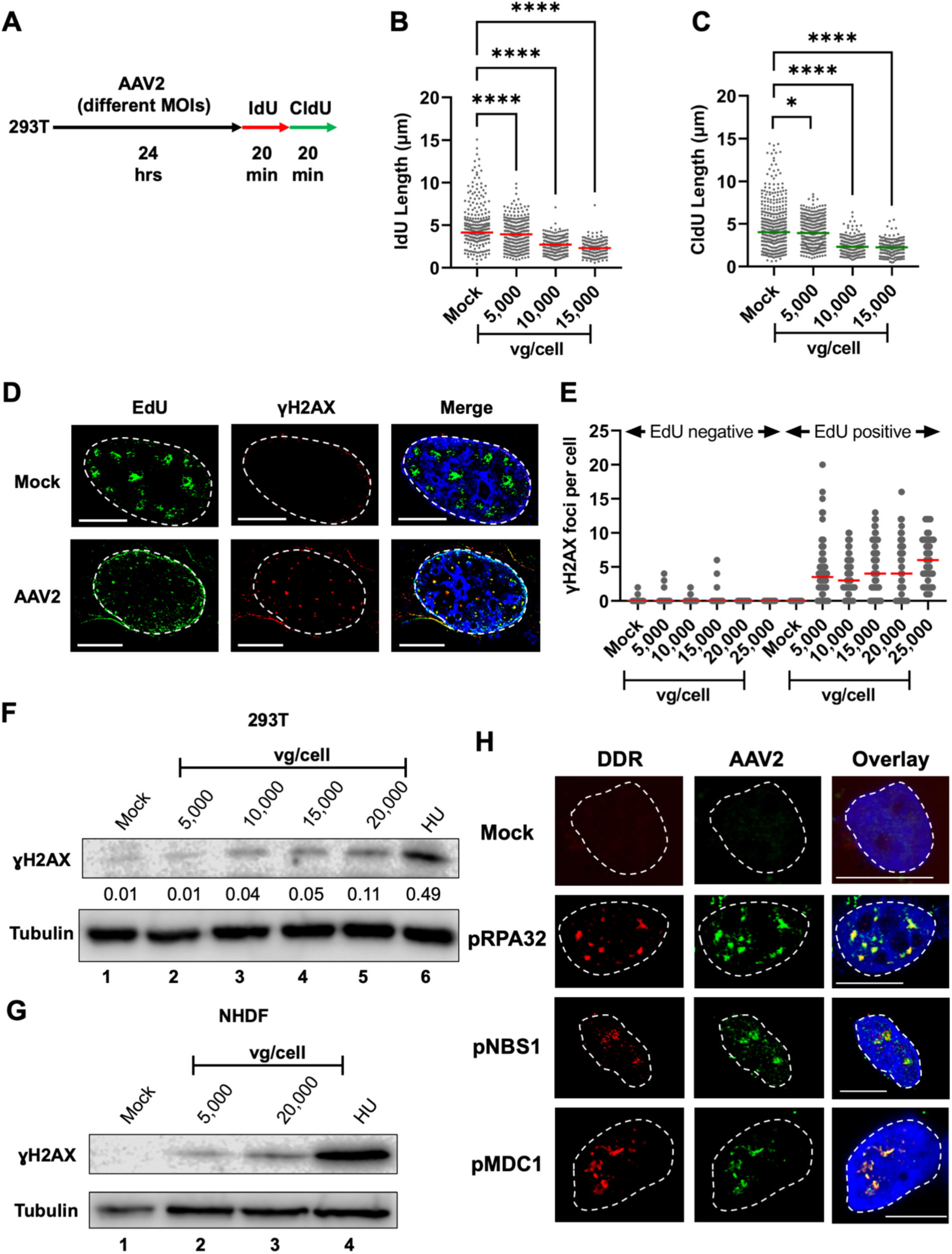
AAV2-mediated RPA32 exhaustion induces cellular DNA damage. (A) Schematic of the timeline of AAV2 infection at different multiplicities in 293T cells for 24 hours before sequential pulsing with IdU and CldU for 20 minutes each prior to DNA fiber analysis. (B,C) DNA fiber length measurements were indicated by each datapoint for (B) IdU-labelled DNA fibers and (C) CldU-labelled DNA fibers. The median lengths of many measurements of IdU and CldU labelled fibers in AAV2 infected cells are represented by red bar and green bar respectively. At least 150 individual fibers were measured for each condition for three independent replicates of AAV2 infections of 293T cells. Statistical significance was measured by Mann Whitney Wilcoxon test, **** represents P < 0.0001, ns represents differences that are not statistically significant. (D,E) U2OS cells infected with AAV2 at an MOI of 10,000 vg/cell were pulsed with EdU (green) for 20 minutes before being processed for gamma H2AX localization (red). The nucleus is shown by DAPI staining (blue) with the nuclear border demarcated by dashed white line. (E) Quantification of the number of gamma H2AX foci per cell in U2OS cells that were EdU-negative (left) and EdU-positive (right) upon being infected with AAV2 at different MOI’s for 24 hours. The median number of gamma H2AX foci in at least 30 nuclei is indicated by red bar. (F,G) Immunoblot analysis of AAV2 infection at the indicated MOI’s compared with 2 mM HU for 24 hours in (F) 293T cells and (G) NHDF cells. Cells were harvested as described and analyzed for γH2AX in the nuclear lysates. Tubulin levels were used as loading controls for the immunoblots. (H) Immuno-FISH assays imaging the relative location of the AAV2 genome (green) to active forms of the identified cellular DDR factors (red) in iPOND-MS studies described above. U2OS cells were infected with AAV2 at an MOI of 10,000 vg/cell for 24 hours before being processed for Immuno-FISH imaging. Nuclei are represented by blue DAPI staining and the nuclear borders are demarcated by dashed white lines. The scale bars represent 10 microns distance.

## DISCUSSION

In this study we have discovered that AAV2 mono-infection as well as rAAV2/scAAV2 transduction at high MOI induces replication stress on the host genome. Since rAAV2/scAAV2 are sufficient to induce replication stress, despite lacking the Rep 68/78 ORFs, these observations suggest that the AAV2 Rep 68/78 proteins are not necessary to induce cellular replication stress. We also observed that neither the ITR element nor empty capsids induced host-cell replication stress. We interpret these results to indicate that the incoming unreplicated single-stranded AAV2 genome is driving replication stress in the infected cells. The observation that AAV2 genomes induce replication stress within to 12 to 24 hours post-infection suggests that acute stress-response signals are likely not involved in shortening cellular DNA synthesis. This is critical because replication stress has recently been connected to innate immune activation upon reactivation of endogenous retroviruses [68]. The absence of further decrease of DNA synthesis at 36 hpi is distinct from that observed in replicating autonomous parvoviruses like MVM where there continues to be a decrease in DNA fiber lengths till cell death [23]. An alternate explanation for this outcome might be that cells that have undergone division within this window have diluted the number of AAV2 genomes inside each cell nucleus, leading to a rescue of RPA exhaustion-mediate stress induction. As a corollary, host-cell RPA molecules associate with AAV2 genomes at high-doses, likely rendering the cellular genome vulnerable to replication stress. Exacerbated by large number of AAV2 genomes in the nucleus, RPA exhaustion causes shortening of cellular replication forks that leads to the induction of cellular DDRs.

Our findings of non-replicative AAV2 genomes inducing replication stress mirror the observations made with UV-inactivated MVM genomes that cause replication stress via RPA exhaustion [23]. Importantly however, while UV-inactivated MVM genomes did not induce cellular γH2AX signals [1], our findings show that AAV2 at high multiplicity did lead to induction of cellular γH2AX. These cellular DDR signals might be generated by virally induced challenges to cell cycle progression. MVM infection inhibits the DREAM complex protein FoxM1 to inhibit transcription of the mitotic entry protein *Ccnb1*, leading to a potent pre-mitotic cell cycle block at the G2/M border [69]. On the other hand, AAV2 infection enriches the transcription factor TFDP2, which interacts with E2F proteins to regulate G1/S checkpoint entry. However, since RNAi-mediated TFDP2 knockdown did not impact AAV2 Rep 68/78 transcripts, our studies suggests that factors like TFDP2 regulate proteins on the host genome and not on the viral genome. While it is important to note that AAV2-induced replication stress does not arrest AAV2-infected cells in S phase, as is the case for parvoviruses like HBoV [70], this replication stress leads to the induction of cellular DNA damage at high multiplicities. These observations suggest that cellular response pathways are likely activated to overcome AAV2-induced replication stress at low multiplicities, allowing for faithful host DNA replication to progress. However, the identity of these host response pathway proteins and how they function remain unknown. We hypothesize that targets identified in our iPOND analysis, such as GINS3, TOP2A and XRCC1, may play a role in maintaining host replisome stability as AAV2 mono-infection progresses long-term.

Recent investigations of iPOND-MS in viral systems have extensively profiled the host factors that are usurped by viruses to amplify their genomes, including polyomaviruses [71], herpesvirses [72–74] and adenoviruses [72, 75]. Since AAV2-monoinfection does not generate nascent DNA at late timepoints, our studies are the first to profile the impact of viral infection exclusively on the host replisome. Indeed, our iPOND-MS analysis point to two distinct possibilities by which viral infection might cause replication stress: 1) formation of protein-DNA complexes, such as those formed by transcription-replication collisions on the host genome; or 2) depletion of replication stress proteins at eukaryotic replisomes by competition from AAV2 genomes. In support of both possibilities, iPOND-MS revealed the enrichment of RNA processing machinery and proteins that regulate replication checkpoints at nascent DNA in AAV2-infected cells. Indeed, the enrichment of PARP1 and CSNK1A2 (Casein Kinase subunits) have previously been found to associate with Rep 68/78 [76] and assembled capsids [77] respectively. CSNK1A2 has been proposed to serve as a host restriction factor that regulates rAAV2 transduction [62]. Additionally, in the MVM system, the non-structural protein NS1 redirects Casein Kinase subunits to suppress local ATR signaling [40]. The details of how ATR-mediated signals may synergize with Casein Kinase to regulate AAV2 life cycle and reduced host DNA synthesis warrants further investigation.

Our iPOND, RNAi and inhibitor studies suggest that RPA plays a key role in AAV2 life cycle and AAV2-induced replication stress. The association of all three RPA subunits (RPA14, RPA32 and RPA70) on the AAV2 genome is striking because RPA canonically binds single-stranded DNA but by 24 hpi most of the AAV2 genomes should have been converted into its double stranded form [63, 78]. It is noteworthy, however, that RPA does bind to double-stranded DNA, albeit at lower binding strength than single-stranded DNA [79]. Alternatively, it is possible that RPA subunits associate with AAV2 genome components that are partially accessible, such as the viral replication origin, polypyrimidine tracks in the ITR or viral promoters. The ChIP-qPCR assays deployed for these studies only sonicate the AAV2 genome to sizes of 500 base pairs or less, making it difficult to resolve the position of the RPA molecules accurately. Regardless, the derepression of AAV2 gene expression upon RPA inhibition indicates that the RPA subunits can directly or indirectly block AAV2 gene expression. One possibility is RPA directly preventing the access of transcription factors. Alternatively, RPA might recruit host factors that transcriptionally silence AAV2 gene expression. An example of this possibility is RPA-mediated recruitment of chromatin modifying enzymes such as G9A [80], a histone methyltransferase that is known to methylate host chromatin [81]. These observations are parallel to the binding of KAP1 to the AAV2 genomes that recruit the chromatin modifier SETDB1 leading to AAV2 gene repression [82]. Future studies will build on how DDR factors on the AAV2 genome generally regulate the chromatin landscape of AAV2 and how this regulates the non-replicative life cycle of AAV2.

Immuno-FISH imaging of AAV2 genomes colocalizing with replication and DDR factors corroborate prior findings of AAV2’s interaction with DDR sites [27]. These nuclear foci are likely sites where the AAV2 genomes undergo processing to be converted from single- to double-stranded DNA molecules, perhaps forming the stalled replication fork-like structures that have previously been described [41]. Since AAV2 infection induces replication stress that can lead to DNA damage, we surmise that AAV2-induced replication stress is focused on cellular sites that are in proximity to the viral genome by usurping RPA molecules from the host. We propose that this competition of RPA molecules renders the host genome vulnerable to replication stress that can lead to DSBs (Fig. 6). These observations are analogous to MVM-induced RPA exhaustion that we have previously discovered [23]. However, unlike replicating MVM genomes, non-replicative AAV2 does not generate multiple single-stranded and double stranded intermediates, rendering the possibility of RPA binding new genome structures unlikely. Taken together, our studies reveal the molecular underpinings of how AAV2-induced replication stress can cause cellular DNA damage.

**Fig. 6:**
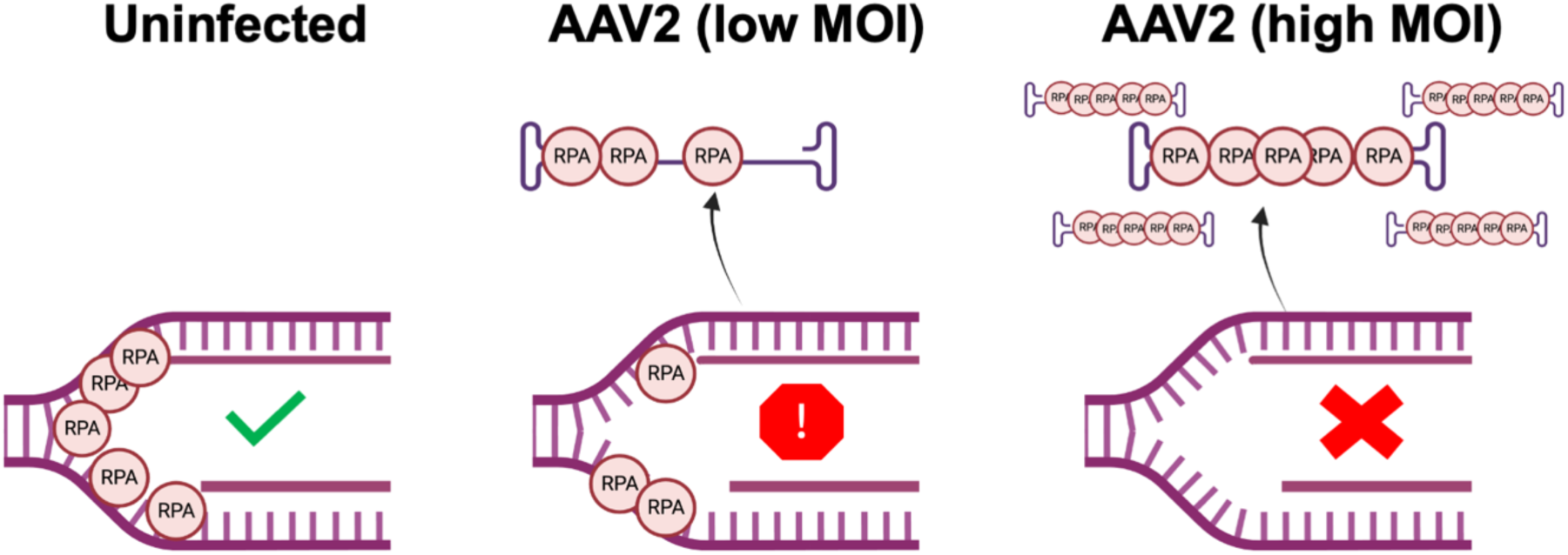
Mechanism of AAV2-induced RPA exhaustion that leads to replication stress on the host genome. AAV2 infection depletes the host genome of RPA molecules, rendering the host replisomes vulnerable to replication stress (middle panel). At high MOI’s of AAV2 infection, this RPA exhaustion leads to induction of cellular DNA damage (right panel). Image generated using Biorender.

## MATERIALS AND METHODS

### Cell Lines

Female human U2OS osteosarcoma cells and female human embryonic kidney cells (HEK293T) were cultured in Dulbecco’s modified Eagle’s medium (DMEM, high glucose; Gibco) supplemented with 5% Serum Plus (Sigma Aldrich) and 50 μg/ml gentamicin (Gibco). Expi293T cells were cultured in Expi293 Expression Medium (Gibco) and were used for AAV2 virus production from the SSV9 infectious clone. Normal Human Dermal Fibroblasts (NHDFs) were obtained from Dr. Robert Kalejta and were cultured in DMEM supplemented with 5% FBS. Cells were cultured in incubators in 5% CO2 at 37 degrees Celsius. Cell lines are tested for mycoplasma contamination and background levels of DNA damage by γH2AX staining.

### Virus and viral infection

AAV2 virus was produced in Expi293T cells and HEK293T cells as described previously [1]. AAV2, scAAV2, and rAAV2 infection and transduction was carried out at a Multiplicity of Transfection (MOT) of 5,000 viral genomes/cell unless otherwise noted. Empty AAV2 capsids were obtained from PROGEN (American Research Products Inc.).

### RNAi, Transfections and Inhibitors

The RNAi sequences (Life Technologies) to silence the respective genes are: RPA2 (GCACCUUCUCAAGCCGAAAtt), TOP2A (*GGAUUCUGCUAGUCCACGAtt*), PRIM1 (*GAACCAGAGAUUAUAAGAAtt*), SIK3 (*CAGCGACGAUGCUUAUGUAtt*), TFDP2 (*GGACUACUUCUGAACUCUAtt*), GINS3 (*GACUUUCAGUGUUGGGAGAtt*), SFN (*ACUUUUCCGUCUUCCACUAtt*), XRCC1 (*GGCAGACACUUACCGAAAAtt*) and CSNK2A1 (*GGCUCGAAUGGGUUCAUCUtt*). RNAi constructs were transfected using Lipofectamine RNAiMAX Transfection kit (Life Technologies) into 293T cells for 24 hours. The RPA inhibitor, TDRL-505 (Millipore), was used at a final concentration of 50 μM in 293T and U2OS cells. For AAV2 virus production, Expi293T cells were transfected with the ExpiFectamine 293 Transfection Kit (Thermo Scientific). The non-palindromic core ITR sequence used to measure ITR-induced replication stress was: ggccgcccgggcaaagcccgggcgtcgggcgacctttggtcgcccggcc (IDT). The RPA over-expression plasmid (pRPA) has previously been published [83] and 0.5 μg was transfected with Lipo293D transfection reagent (SignaGen). An equivalent amount of pUC18 plasmid was used as mock transfection control.

### DNA Fiber Assay (DFA)

HEK293T or U2OS cells were transduced or infected according to experimental requirements (as described above and previously [84]). Single molecule DNA Fiber Analysis experiments were completed following previously published protocols [58]. Cells were pulsed in 20mM IdU for 20 minutes at the end of infection, followed immediately by pulsing with 50mM CldU for 20 minutes. Cells were pelleted at 5000xg for 5 minutes and resuspended in 200 μL of complete media, then stored on ice for the duration of the protocol. 2 μL of resuspended cell solution was pipetted onto positively charged slides, then mixed with 6 μL of DFA Lysis Buffer (200 mM Tris-HCl pH 7.5, 50 mM EDTA, 0.5% SDS) and allowed to lyse for 5 minutes. Slides were then tilted to spread the DNA fibers along the slide and air dried for 15 minutes. DNA was fixed using a 3:1 methanol:acetic acid solution. The DNA was denatured using 2.5 M HCl for 1 hour at room temperature. After denaturing, slides were blocked in 3% BSA in PBS for 30 minutes in a humidified chamber at room temperature. Primary antibody staining was carried out using Abcam rat anti-BrdU (1:1000) and BD Biosciences mouse anti-BrdU (1:500) at room temperature for 30 minutes, then washed with 0.1% Tween 20 in PBS 3 times. Slides were then stained with anti-rat Alexa Fluor 488 and anti-mouse IgG1 Alexa Fluor 568 (1:1000) secondary antibodies at room temperature for 30 minutes in the dark. Samples were washed with 0.1% Tween 20 in PBS 3 times and cover slips were affixed to slides using ProLong Gold Antifade Mountant (Thermo Scientific). Fibers were imaged with a Leica Stellaris confocal microscope using a 63X oil immersion objective lens. Fiber lengths were measured using Digimizer software (MedCalc Software Ltd).

### Antibodies

Antibodies used for DNA fiber analysis were: anti-BrdU (BD Biosciences, Clone B44, 347580), anti-BrdU (Abcam, ab6326), Alexa-Fluor 568 conjugated anti-mouse secondary (Thermo Scientific, A11004), Alexa-Fluor 488 conjugated anti-rat secondary (Thermo Scientific, A11006).

Antibodies used for western blot: γH2AX (AbCAM, ab11174), Tubulin (Millipore, 05-829); ChIP-qPCR analysis: RPA14 (Life Technologies, MA1-23281), RPA32 (Cell Signaling, 52448S), RPA70 (Cell Signaling, 2267S) and phospho-RPA32 Ser 8 (Cell Signaling, 83745); Immuno-FISH: phospho-MDC (AbCAM, ab36513) and phospho-NBS1 (Cell Signaling, 3001).

### Chromatin Immunoprecipitation combined with Quantitative PCR (ChIP-qPCR)

HEK293T cells were infected with AAV2 as described above and were crosslinked in 1% Formaldehyde for 10 minutes at room temperature. 0.125 M glycine was used to quench the crosslinking reaction for 5 minutes at room temperature. Cells were lysed on ice for 20 minutes in ChIP lysis buffer (1% SDS, 10 mM EDTA, 50 mM Tris-HCl, pH 8, protease inhibitor). Cell lysates were sonicated using a Diagenode Bioruptor Pico for 60 cycles (30 s on and 30 s off per cycle), before being incubated overnight at 4 degrees C with the antibodies bound to Protein A Dynabeads (Invitrogen). Using low-salt wash (0.01% SDS, 1% Triton X-100, 2 mM EDTA, 20 mM Tris-HCl pH8, 150 mM NaCl), high salt wash (0.01% SDS, 1% Triton X-100, 2 mM EDTA 20 mM Tris-HCl pH8, 500 mM NaCl), lithium chloride wash (0.25M LiCl, 1% NP40, 1% DOC, 1 mM EDTA, 10 mM Tris HCl pH8) and twice with TE buffer samples were washed for 3 minutes each at 4 degrees. Using SDS elution buffer (1% SDS, 0.1M sodium bicarbonate) DNA was eluted and crosslinks were reversed using 0.2M NaCl, Proteinase K (NEB) and incubated at 56 degrees C overnight. Using a PCR Purification Kit (Qiagen) DNA was purified and eluted in 100 μl of Buffer EB (Qiagen). ChIP DNA was quantified by qPCR analysis (Biorad) under the following conditions: 95^0^ C for 5 mins, 95^0^ C for 10 secs and 60^0^ C for 30 secs for 40 cycles. AAV2 genome interaction with RPA molecules was assessed by qPCR assays using primers complementary to the REP open reading frame. These values were compared with primers monitoring the levels of Beta actin transcripts. The primer sequences used for RT-qPCR and ChIP-qPCR on the Rep 68/78 ORF in 5’ to 3’ orientation are: TGATAAGCGGTTCAGGGAGT (forward primer) and CCAGCCATGGTTAGTTGGTT (reverse primer); Beta actin gene: CACCTTGATCTTCATTGTGCTG (forward primer) and GCAAAGACCTGTACGCCAAC (reverse primer).

### Statistical analysis

Statistical analysis was performed using Graphpad Prism software and appropriate statistical test as indicated in the manuscript.

### iPOND-SILAC-MS

iPOND was performed as previously described [61]. Cells (U2OS or 293T) were labeled with 10μM EdU for 10 minutes following 24-hour infection at an MOI of 5,000 vg/cell with AAV2 virus. Cells were cross-linked in 1% fresh formaldehyde/PBS for 10 minutes (RT), quenched with 1.25 M glycine, and washed with PBS. Collected cell pellets were frozen at -80 C. The next day pellets were permeabilized in 0.25% Triton-X/PBS and washed once with 0.5% BSA/PBS followed by PBS. Light and heavy labeled cells were mixed 1:1 by cell number and the click reaction was completed in 1 hour with PEG4-biotin azide (Invitrogen). The cells were subsequently lysed by sonication using a Diagneode Pico sonicator on ultra-high setting until homogenized. DNA-protein complexes were purified using streptavidin-coupled C1 magnetic beads for 1 hr. Samples were washed (5 minutes each) with lysis buffer (1% SDS in 50 mM Tris, pH 8.0), low salt buffer (1% Triton X-100, 20 mM Tris, pH 8.0, 2 mM EDTA, pH 8.0, 150 mM NaCl), high salt buffer(1% Triton X-100, 20 mM Tris, pH 8.0, 2 mM EDTA, pH 8.0, 500 mM NaCl), lithium chloride buffer (100 mM Tris, pH 8.0, 500 mM LiCl, 1% Igepal), followed by two washes in lysis buffer. Captured proteins were eluted and cross-links were reversed in 6X SDS sample for 30 min at 95 C while mixing every 10 minutes.

All incubations were carried out at room temperature unless otherwise specified. iPOND eluates were separated on 4–12% Tris-glycine mini gels (Novex™, Invitrogen). Following electrophoresis, gels were incubated for 30 min in fixing solution (50% methanol, 1.2% phosphoric acid) to immobilize proteins. After a brief rinse in water, gels were stained overnight in a solution containing 50% methanol, 1.2% phosphoric acid, 1.3 M ammonium sulfate, and 0.1% (w/v) Coomassie Brilliant Blue G-250.

Each lane (sample) was then cut into small rectangular pieces and divided into two fractions: fraction 2 (containing the streptavidin band) and fraction 1 (containing the remaining proteins). For in-gel digestion, both fractions were subjected to several rounds of washing to remove salts and Coomassie. Specifically, gel pieces were incubated for 15 min in 25 mM ammonium bicarbonate, 50% ethanol (60 °C for strongly stained pieces), and the supernatant was discarded. This wash step was repeated 2–3 times or until gel pieces were destained completely. Next, in preparation for cysteine reduction & alkylation, fractions were incubated in 100% ethanol for 10 min to dehydrate the gel pieces.

Cysteine residues were reduced by incubating fractions for 20 min at room temperature in 20 mM DTT, 50 mM ammonium bicarbonate. After discarding the supernatant, cysteine residues were alkylated by a 20-min incubation in 80 mM chloroacetamide, 50 mM ammonium bicarbonate. Alkylation compounds were removed with successive 10-min washes in i) 50 mM ammonium bicarbonate, ii) 25 mM ammonium bicarbonate, 50% ethanol, and iii) 100% ethanol. Gel pieces were subsequently dried in a vacuum concentrator.

For overnight digestion at 37 °C, trypsin solution (0.25 µg trypsin [Promega] in 50 µl of 50 mM ammonium bicarbonate) was added to each fraction. On the following day, peptides were extracted by two sequential 15-min elutions in 150 µl of elution solution (80% acetonitrile, 0.2% formic acid) under sonication. The resulting eluates were pooled and evaporated to dryness in a vacuum concentrator.

Prior to LC–MS analysis, dried peptide samples were subjected to a final cleanup using solid-phase extraction (SPE) cartridges (1 cc, 50 mg C18 sorbent, Sep-Pak, Waters) in conjunction with a vacuum manifold. Samples were first resuspended in 200 µl of 0.2% formic acid. The cartridges were then equilibrated with 100% acetonitrile, followed by two washes with 0.2% formic acid. After loading the samples onto the cartridges, two additional washes with 0.2% formic acid were performed. Finally, peptides were eluted with 80% acetonitrile, 0.2% formic acid, and the eluates were dried to completion in a vacuum concentrator. Prior to LC–MS analysis, dried peptide samples were resuspended in 0.2% formic acid and their concentrations were determined using a NanoDrop spectrophotometer.

### LC-MS analysis

For LC–MS analysis, a Vanquish Neo UHPLC system (Thermo Fisher Scientific) was coupled to an Orbitrap Ascend mass spectrometer (Thermo Fisher Scientific) via a Nanospray Flex ionization source. A 40 cm in-house–prepared fused silica C18 column was maintained at 50 °C using a custom-built column heater. The source voltage was set to 2 kV, and the ion transfer tube was held at 275 °C. Peptide samples from streptavidin-containing fractions were injected at a total amount of 500 ng, whereas 1 µg of peptides was loaded for all other samples. Peptides were separated at a flow rate of 300 nL/min with a 73 min active gradient from 5% to 46% solvent B (solvent A: 0.2% FA; solvent B: 80% acetonitrile with 0.2% FA).

Orbitrap Ascend parameters were as follows: MS1 scans were acquired at a resolution of 240k (scan range 300–1350 m/z), with a maximum injection time of 50 ms, an automatic gain control (AGC) target of 1 × 10^6^, a normalized AGC target of 250%, and an RF lens percentage of 30. Data-dependent acquisition employed a 1 s cycle time. MIPS (monoisotopic peak determination) was set to “peptide”, and the isolation window center was set to “most abundant peak”. Charge states 2–5 were included, and dynamic exclusion was set to 20 s with a ±5 ppm mass tolerance. MS2 spectra were recorded in the ion trap using an isolation window of 0.8 m/z, a normalized HCD collision energy of 24%, an ion trap scan rate set to “Turbo,” a scan range of 150–1350 m/z, a normalized AGC target of 250%, and a maximum injection time of 12 ms.

### Proteomics data analysis

RAW files generated from the LC–MS runs were analyzed with MaxQuant (version 2.4.2.0), searching against a UniProt human FASTA file containing both SwissProt and TrEMBL entries. Adeno-associated virus 2 proteins (Rep68 [P03132] and Rep78 [Q89268]) were added to this database. Unless otherwise specified, all MaxQuant parameters were set to their default values.

Gel fractions originating from the same lane were assigned identical experiment names but treated as separate fractions in MaxQuant. In the *Group-specific parameters* tab, “Type” was set to *multiplicity = 2*, and *Arg10* and *Lys8* were checked. Under *Misc.*, *Re-quantify* was enabled. In the *Global parameters* tab, the FASTA files described above were specified under *Sequences*. Under *Protein quantification*, *Label min. ratio count* was set to *1*, and under *Identification*, *Min. unique peptides* was set to *1* with *Match between runs* enabled. Finally, under *Label-free quantification*, *iBAQ* was enabled.

R (v4.4.3) was used to visualize mass spectrometry–derived gene expression data, integrating gene ontology (GO) annotations and essential biological pathways. Data were imported with readr, and biomart annotations retrieved via biomartr. Gene IDs were mapped to GO terms and pathways using org.Hs.eg.db, followed by functional enrichment analysis with clusterProfiler. High-quality, customizable plots were generated in ggplot2, depicting enriched GO terms and pathway networks. Analyses were conducted in RStudio to ensure reproducibility and modular code.

### AAV2 Virus production

Expi293F cells were passaged to a cell density of 2.0×10^6^ – 3.0×10^6^ cells/mL resuspended in 25 mL of fresh Expi293F media and expected to double within 24 hours when incubated at 37 degrees C, 8% CO2 in a shaking incubator. To prepare for transfection, cell culture volumes were expanded to 1L at the same density, then diluted according to manufacturer’s specifications. PEI (1 mg/mL concentration) was used as the transfection method at a ratio of 3:1. Plasmid DNA (1μg/mL of cell culture) and PEI were diluted in OPTI-MEM media in separate tubes according to manufacturer’s guidelines. These mixtures were combined and incubated for 10-15 minutes at room temperature to generate transfection complexes. The complexed mixtures were added to the cell culture dropwise as the flask was being swirled. The next day of transfection, enhancer 1 and 2 solutions were added from the Expifectamine Transfection kit (Thermo Scientific) and placed back into the incubator for the remainder of the incubation. After 5-7 days of incubation the cells were collected into 50 mL falcon tubes and centrifuged for 2 minutes at maximum speed. The pellet was resuspended in 15 mL of TE buffer. A series of seven freeze-thaw cycles were performed by alternating between liquid nitrogen (for freezing) and 37^0^C incubator (for thawing). After the freeze/thaw cycles DNase I (Thermo Scientific) treatment was carried out with 0.5U/μL for 1 hour at 37^0^C. The cells were then centrifuged at 2,000xg at 4^0^C for 10 minutes and the supernatant was transferred into a screw cap tube and stored in 4^0^C for temporary storage, and -80^0^C for long-term storage.

### Immuno-FISH imaging

DNA probe was generated by labelling oligonucleotides complementary to the AAV2 genome using Aminoallyl-1-dUTP and TdT (Thermo Scientific). The oligos were ethanol precipitated and NHS-ester dyes (Thermo Scientific) were conjugated using sodium bicarbonate. Probes were purified using PCR clean-up column (Promega) and dissolved in TE buffer, as previously described [85]. After 16-24 hours post infection, cells were pre-extracted in Cytoskeletal (CSK) buffer for 3 minutes followed by 1 mL of CSK+ Triton X solution for 3 minutes. CSK buffers were aspirated, cells washed with 1 mL PBS and fixed with 4% paraformaldehyde in PBS for 10 min at room temperature (RT). Cells were washed with 1 mL of PBS and permeabilized in 1 mL of Permeabilization Buffer for 10 minutes. 10% formamide in 2X SSC buffer (300 mM sodium chloride, 30 mM sodium citrate) was added to the cells for 1-2 hours to rehydrate and pre-denature the DNAs. The probes were suspended in hybridization buffer to a final concentration of 0.5 ng/μL. Samples were hybridized with the probe solution in 2XSSC and sealed on glass slide with rubber cement and hybridized overnight at 37 degrees C in a humidified chamber. Cover slips were washed in 2X SSC/0.1% Triton X-100 at 37 degrees for 3 minutes each, 2X SSC at 37 degrees C and mounted on slide with Prolong Diamond Antifade Mounting Media with DAPI. Samples were imaged on a confocal microscope (Leica) with 63X oil objective lens.

### EdU labeling coupled with Immunofluorescence Imaging

U2OS cells were plated on coverslips in 6-well plates and allowed to adhere overnight before being infected with AAV2 at the indicated MOIs for 24 hours. EdU labelling was carried out using 10 mM EdU stock solution diluted to a final concentration of 20 μM. The cells were pulsed with EdU for 2 hours. After the 2-hour incubation the samples were fixed using 4% PFA for 15 minutes at room temperature. Then the cells were washed with PBS and permeabilized with 0.5% Triton X-100 for 20 minutes at room temperature. After the incubation wash with PBS, 500 μL of the Click-it Reaction Cocktail (containing 1X Click-iT EdU reaction buffer, Copper sulfate, Alexa-Fluor-488 Azide and EdU buffer additive) was added to each sample and incubated for 30 minutes. Samples were washed with PBS. The samples were stained using 3% BSA in PBS (blocking) followed by incubation with the primary antibody for 30 minutes at room temperature. Samples were washed with PBS. Samples were stained with fluorophore conjugated secondary antibody in 3% BSA in PBS for 30 minutes in the dark at room temperature. Samples were washed in PBS mounted onto coverslips using DAPI-containing Fluromount. The samples were imaged on a Leica confocal microscope with 63X oil immersion objective.

### RNA extraction

Cells were plated and infected for 24 hours. The cells were then harvested and resuspended in 1 mL of PureZOL (Bio-Rad). 200 μL of Chloroform was then added to the microcentrifuge tubes and shaken vigorously for 15 seconds each. The samples were then incubated for 5 minutes at room temperature on a rotator. The samples were centrifuged at 12,000xg for 15 minutes at 4^0^C. The aqueous layer was collected and transferred into a new microcentrifuge tube. The cells were precipitated with 2 μL of glycogen and 500 μL of isopropyl alcohol at minus 20 degrees C. Samples were centrifuged at 12,000xg for 10 minutes in 4^0^C. The RNA pellet was washed in 1 mL of 75% ethanol, centrifuged at maximum speed for 5 minutes at 4^0^C and air dried for 5 minutes. After the pellet was dried DNase treatment (Promega) was performed and incubated for 30 minutes at 37^0^C. The samples were then incubated at 65^0^C for 10 minutes to inactivate DNase. DNase stop solution was added to the sample. The RNA pellet was resuspended in PureZOL (Bio-Rad) and the purification was repeated. The pellet was resuspended in 20 μL of water. Reverse Transcription (RT) reaction was carried on 1 μg of RNA using the iScript cDNA synthesis kit (Bio-Rad). The generated cDNA was diluted in 80 μL of water.

### Western blots

Cells were plated in 6 well plates at 500,000 cells and infected with AAV2 virus for 24 hours. After the 24 hours cells were harvested and processed by lysis in RIPA buffer (containing protease inhibitors, sodium orthovanadate and sodium fluoride) on ice for 15 minutes. The cells were centrifuged at 13,000 rpm for 10-minutes at 4^0^C, supernatants collected and denatured with 6X Loading dye. Approximately 40 micrograms of lysate [measured using BCA assay (Bio-Rad)] were loaded on SDS-PAGE gels at 80-150V. Samples were transferred to a PVDF membrane using semi-dry transfer conditions at 25V for 30 minutes. The membrane was blocked with 5% milk in TBST for 30 minutes, incubated with appropriate primary antibody (1:1000, 1:2500, or 1:5000 dilution) and relevant secondary antibody (1:5000 dilution) for 1 hour each. Samples were incubated with ECL reagent (Bio-Rad) for 5 minutes and imaged on the LiCOR instrument.

## ACKNOWLEDGEMENTS

We thank all members of the Majumder Lab for insightful discussions and Rhiannon R. Abrahams (Majumder Lab, UW Madison) for expert technical guidance in generating AAV2 viruses. We thank Dr. David Pintel (University of Missouri) for scAAV2-GFP vector and Dr. Paul Lambert (UW Madison) for critical reading of the manuscript. This research was funded partially by NIH/NIAID K99/R00 Pathway to Independence Award, grant number AI148511 to K.M.; Wisconsin Partnership Program’s New Investigator Award (PERC Grant G-4942) to K.M.; NIH/NIGMS R35 Maximizing Investigator’s Research Award (MIRA), grant number GM154938 to K.M and NIH P41GM108538 to J.C. C.I.S.L. is funded by an NSF Graduate Research Fellowship Program award 2137424 and M.K.H by an NSF Graduate Research Fellowship Program award 2023348714. M.F.S. is funded by a SciMED Graduate Research Scholarship from the University of Wisconsin-Madison and Molecular and Cellular Pharmacology T32 training grant T32GM141013 from the NIH. The content is solely the responsibility of the authors and does not necessarily represent the official views of the National Institutes of Health. The authors declare no competing financial interests.

